# Structure of Importin-4 bound to the H3-H4·ASF1 histone·histone chaperone complex

**DOI:** 10.1101/2022.04.08.487665

**Authors:** Natalia Elisa Bernardes, Ho Yee Joyce Fung, Yang Li, Zhe Chen, Yuh Min Chook

## Abstract

Importin-4 is the primary nuclear import receptor of core histones H3 and H4. Importin-4 binds the H3-H4 dimer and histone-chaperone ASF1 prior to nuclear import, but available structures of Importin-4·histone tail complexes do not explain how Importin-4 recognizes the biologically relevant heterotrimeric H3-H4·ASF1 cargo. Our 3.5 Å Importin-4·H3-H4·ASF1 cryo-electron microscopy structure revealed interactions with H3-H4·ASF1 different those suggested by previous Importin-H3 tail peptide structures. The N-terminal half of Importin-4 clamps the globular histone domain and the H3 αN helix while its C-terminal half binds the H3 N-terminal tail weakly, with negligible tail contribution to binding energy; ASF1 binds H3-H4 without contacting Importin-4. Together, ASF1 and Importin-4 shield nucleosomal interfaces of H3-H4 to chaperone and import it into the nucleus, where Importin-4 undergoes large conformational changes as RanGTP binds to release H3-H4·ASF1. This work explains the mechanisms of nuclear import of full-length H3-H4.

---

During the S-phase of the cell cycle, core histones H3, H4, H2A and H2B are rapidly synthesized in the cytoplasm, transported into the nucleus and then incorporated into newly replicated DNA to form nucleosomes. Following translation in the cytoplasm, H3 and H4 are passed to heat-shock proteins and histone-chaperones to be folded into H3-H4 heterodimers, which are then acetylated at several lysine side chains (Benson et al., 2006; Campos et al., 2010; Tagami et al., 2004). Multiple studies have shown that H3 and H4 are actively transported across the nuclear pore complex (NPC) into the nucleus by Karyopherin-β nuclear import receptors or Importins (Baake et al., 2001; Muhlhausser et al., 2001; Schwamborn et al., 1998). The major Importins that co-purify with H3 and H4 from *S. cerevisiae* (*Sc*) and HeLa cell extracts are *Sc* KAP123 and human Importin-4 (IMP4), respectively, while smaller amounts of *Sc* KAP121 and human Importin-5 (IMP5) associate with H3 and H4 as secondary or backup Importins (Alvarez et al., 2011; Campos et al., 2010; Jasencakova et al., 2010; Pardal et al., 2019).

Early studies showed that the basic N-terminal tail peptides of both H3 and H4 (H3^tail^ and H4^tail^, Figure 1A) can bind and be imported by at least six different Importins (Mosammaparast et al., 2002; Muhlhausser et al., 2001; Soniat et al., 2016) but nuclear import of the full-length histones appears more complex. For example, the minor Importins for H3 and H4 (Kapβ2, IMP9, IMP5) bind histone tails more tightly than the major/primary importin, IMP4 (Soniat et al., 2016). Crystal structures of the *Sc* homolog of IMP4, KAP123, bound separately to the H3^tail^ (residues 1-28) and H4^tail^ (residues 1-34) showed interactions with only a few histone residues in each structure (An et al., 2017), and cellular studies of H3 and H4 without N-terminal tails suggested that tail-less histones are imported into the nucleus in a manner dependent on their histone-fold domain (Apta-Smith et al., 2018; Blackwell et al., 2007). Most importantly, H3-H4 heterodimers were detected in the cytoplasm of HeLa cells in complex with both IMP4 and the histone-chaperone ASF1, suggesting that the cargo transported by IMP4 is the H3-H4·ASF1 complex (Campos et al., 2010). Of several importins tested, IMP4 also binds the tightest to the H3-H4·ASF1 complex, consistent with its role as the major H3 and H4 importer (Soniat et al., 2016). In the absence of an atomic resolution structure of IMP4 bound to H3-H4·ASF1, it was unclear how the H3-H4 globular histone-fold domain, the histone tails and ASF1 contribute to the formation of the IMP4·H3-H4·ASF1 import complex.

**Figure 1.**
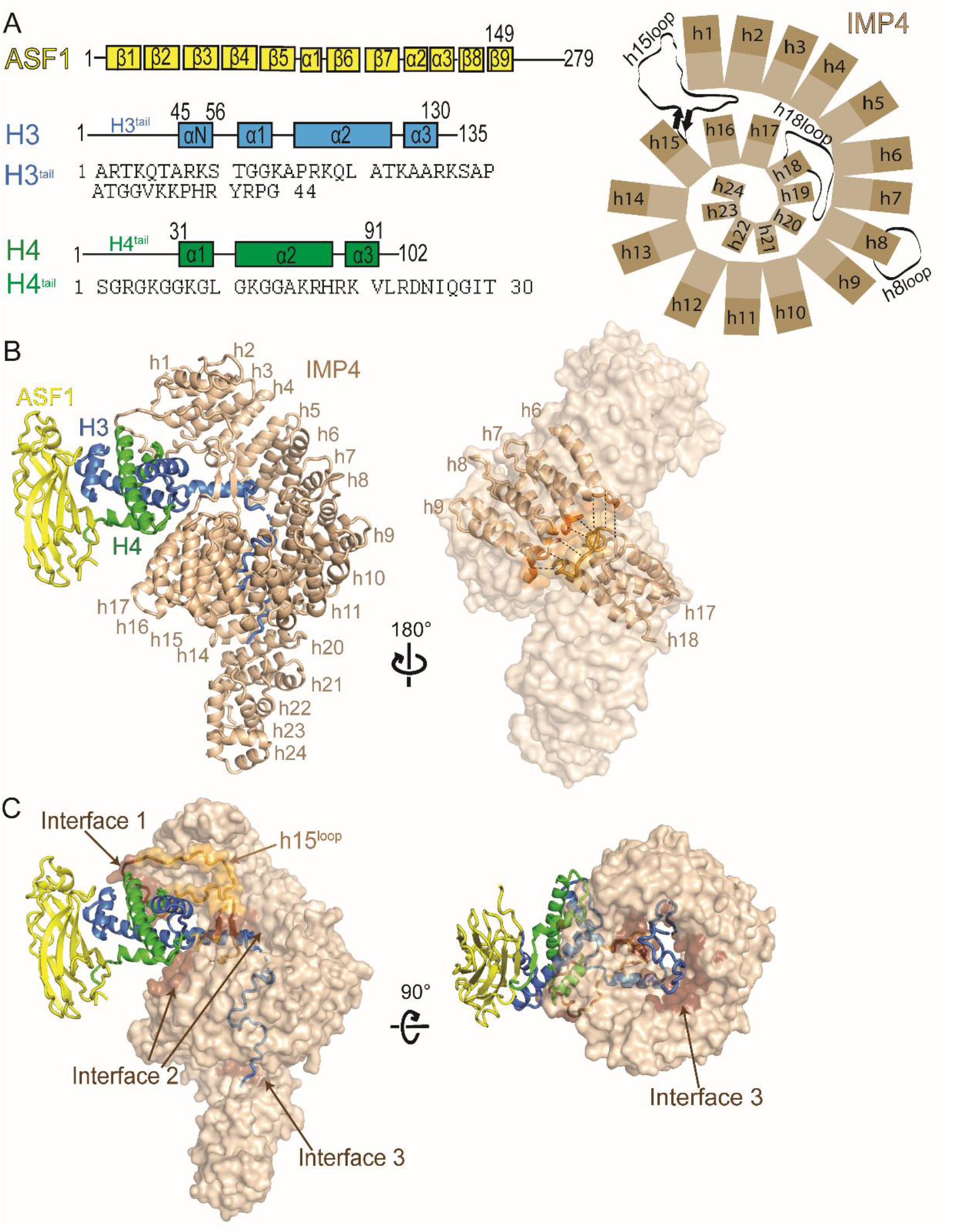
The cryo-EM structure of IMP4·H3-H4·ASF1. A) A schematic showing the organizations of *H. sapiens* IMP4, *S. cerevisiae* ASF1, *X. laevis* H3 and H4 and the sequence of the H3 and H4 N-terminal tails. B) Left, a cartoon representation of IMP4 (beige) bound to H3-H4·Asf1 (H3 is blue, H4 green and ASF1 yellow), with IMP4 HEAT repeats h1-h24 labeled. Right, IMP4 in surface representation, highlighting the two regions, h6-h9 (dark orange) and h17-18 (dark yellow), that interact to close the central ring of the superhelix. C) IMP4 (beige surface) contacts H3-H4 at interfaces 1, 2 and 3 (dark brown surface) with the long IMP4 h15^loop^ colored light orange. A 90° rotation about the horizontal axis shows the tight central ring of IMP4 in the right panel. HEAT repeats h23-h24 are not shown for a clear view of the ring.

We assembled full-length human IMP4, *X. laevis* H3-H4 and *Sc* ASF1 (residues 1-160) to form a 1:1:1 stoichiometry IMP4·H3-H4·ASF1 complex and solved its structure to 3.5 Å resolution by cryo-electron microscopy (cryo-EM) (Figure 1A-C, Figure S1A, B and Table S1). The structure shows one IMP4 molecule bound to one H3-H4·ASF1 complex; the latter is almost identical to a previous H3-H4·ASF1 structure (PDBID:2HUE, rmsd: 0.607 for 243 Cα atoms) (English et al., 2006). IMP4 makes extensive interactions with H3-H4: the globular H3-H4 histone-fold domain and the αN helix of H3 bind to the N-terminal half of IMP4 while the H3^tail^ binds to the C-terminal half of the importin (Figure 1B, C). ASF1 binds the H3-H4 dimer but makes no contact with IMP4. No density is observed for the H4^tail^ and the C-terminal tails of the histones.

Histone-bound IMP4 is an elongated superhelix of 24 HEAT repeats (h1-h24), each comprising a pair of A and B α-helices (Figures 1A, B). Like its yeast homolog Kap123 (An et al., 2017), most IMP4 helices are connected by short loops except for the helices in h8, h15 and h18. The h8^loop^ is 26 residues long and partially disordered (Figure 1A, B). The h15^loop^ is 50-residue long (residues 618-668), contains a β-strand pair at its proximal end and has a highly acidic distal portion. This loop extends from the middle of the IMP4 superhelix towards the IMP4 N-terminus where it interacts with repeats h1-h3 and with the histone-fold domain of H3-H4 (Figure 1A-C, S2A). Interestingly, the h15^loop^ in the unliganded and H3^tail^ peptide-bound Kap123 structures (PDBID: 5VE8 and 5VCH) are disordered and not modeled (An et al., 2017). The highly acidic h18^loop^ (^812^DTDEEEEEEADDQAEYD^828^) of H3-H4·ASF1-bound IMP4 is also involved in intramolecular interactions, making electrostatic interactions with a series of lysine side chains in the h7 repeat (Figure 1A, B, S2B). These h18^loop^-h7 interactions and additional interactions between h17-h18 with h6-h9, bring together the two distant sets of HEAT repeats to form a small ring (inner diameter ~20 Å) in the middle of the IMP4 superhelix that the H3^tail^ threads through (Figure 1B, C). The H3^tail^-bound and the unliganded KAP123 superhelices are similar, with repeats h17-h18 and h6-h9 in close proximity (Figure S2C).

IMP4 makes extensive interactions with H3-H4 through three noncontiguous interfaces 1-3 that cover a total surface area of 1969 Å^2^ on the histone dimer (Figure 1C). The H3-H4 histone-fold domain is clamped by the N-terminal half of the IMP4 superhelix, through interactions at Interface 1 with the distal part of the IMP4 h15^loop^ (which also interacts with IMP4 repeats h1-h2) and with IMP4 h14-h16 (part of Interface 2) (Figure 2A). h15^loop^ residues^636^DDESDGEEEEELMDED^651^ interact with helix α1 of H4 and the loop that follows (loop 1, Figure 2B). Interactions here involve many main-chain contacts, including hydrogen bonds, that resemble β-sheet interactions. Many acidic h15^loop^ side chains make electrostatic interactions with basic H4 side chains; hydrophobic interactions are also present. On the other side of the histone-fold domain, residues in the α2 helix of H3 and residues in the region that span α2-α3 of H4 interact with IMP4 repeats h14-h15 through a myriad of interaction types to form part of Interface 2 (Figure 2C). The other part of Interface 2 involves interactions of the H3 αN helix (αN^H3^, residues 45-58).

**Figure 2.**
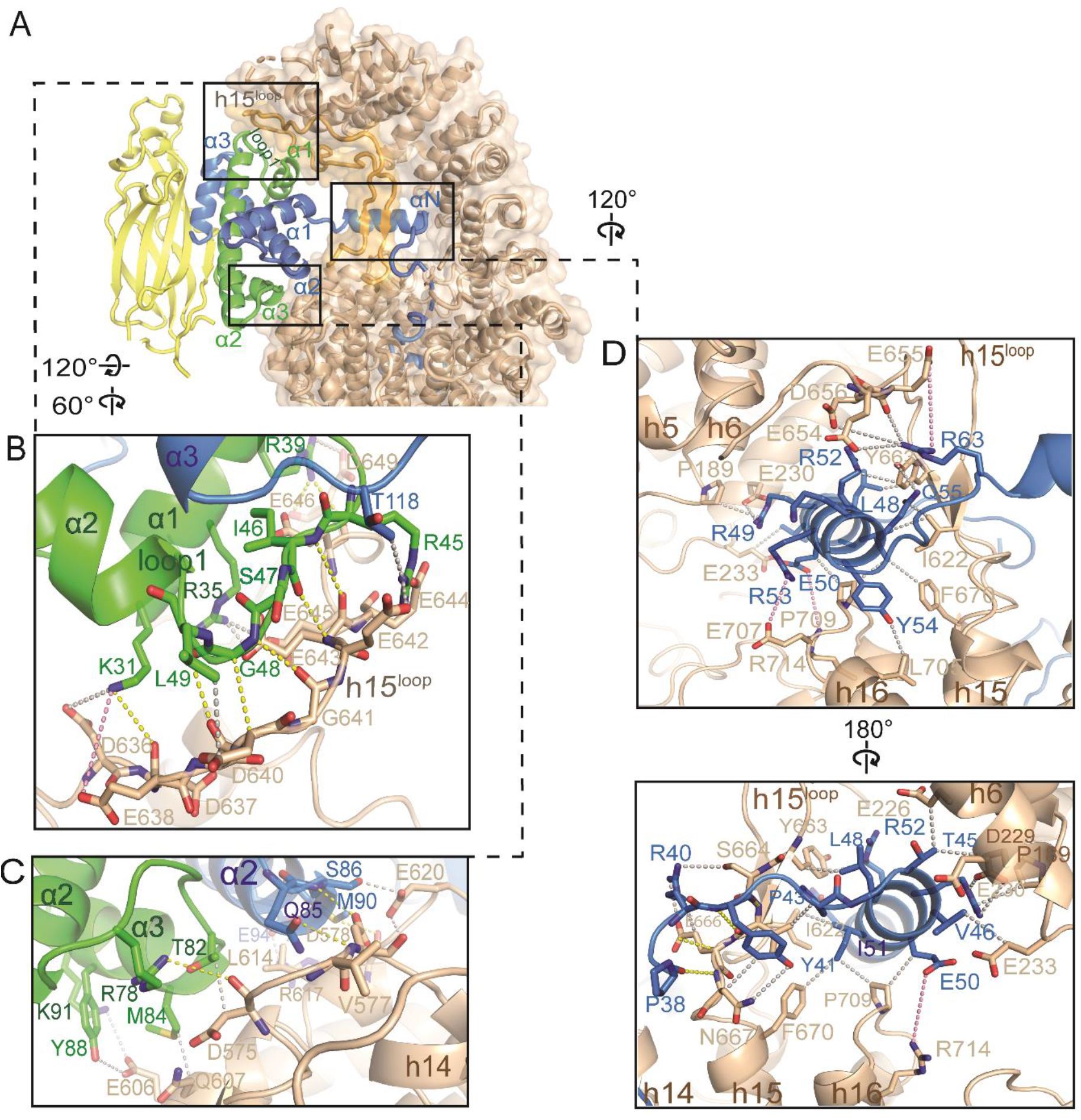
The histone-fold domain of H3-H4 and the αN^H3^ helix binds the N-terminal half of IMP4. A) Cartoon and surface representations of the N-terminal half of IMP4 (beige) clamping the histone-fold domain of the ASF1-bound H3-H4 (ASF1 is yellow, H3 blue and H4 green), interacting at Interfaces 1 and 2. The IMP4 h15^loop^ is colored light orange. B-D) Details of Interfaces 1-2. Select interactions are shown with yellow dashed lines (interactions that involve main chains), gray dashes (interactions between sidechains) and pink dashes (long-range electrostatic interactions). B) Details of Interface 1, between IMP4 h15^loop^ residues and the region of H4 that spans α1-α2. C) Details of Interface 2, between residues of IMP4 HEAT repeats h14 and h15 with the other end of the H3-H4 domain from B), involving residues from α2-α3 of H4 and from the α2 helix of H3. D) Details of the remaining portion of Interface 2, where the proximal part of the h15^loop^, HEAT repeats h14-h16 and h4-h6 of IMP4 surround the αN^H3^ helix.

αN^H3^ adopts different orientations relative to the histone-fold domain depending on the binding partners. αN^H3^ in the nucleosome interacts with and is part of the histone-fold domain as it also makes multiple contacts with DNA whereas αN^H3^ in the H3-H4·ASF1 crystal structure is likely highly dynamic as hence not modeled (English et al., 2006; Luger et al., 1997). However, the αN^H3^ helix in our IMP4·H3-H4·ASF1 structure protrudes away from the histone-fold domain and is surrounded by several different IMP4 elements – B helices of IMP4 repeats h4-h7, the IMP4 h16A helix and the β-strand pair of the IMP4 h15^loop^ – that make many electrostatic, polar and hydrophobic interactions with the H3 helix (Figure 2D).

The segment N-terminal of αN^H3^ (residues 37-44) approach the IMP4 central ring and no density is observed N-terminal of that for H3 residues 26-36 (Figure 3A, S3A-B). At the very N-terminus, H3^tail^ residues 1-26 bind to B helices in the C-terminal half of IMP4 in a direction anti-parallel to the IMP4 HEAT repeat progression through a variety of interactions - electrostatic, polar and hydrophobic (Figure 3A-D). The positioning of the αN^H3^ helix at the opening of the IMP4 central ring threads the H3^tail^ into and through the ring, placing the extreme N-terminus of H3 to bind the C-terminal HEAT repeats of IMP4. H3^tail^ segment ^1^ARTKQ^5^ binds C-terminal HEAT repeats h18-h22, while H3^tail^ residues 7-25 line the inner surface of the IMP4 central ring (Interface 3; Figure 3A-D). Within the IMP4 ring, the H3^tail^ segment ^7^ARKS^10^ contacts h17-h18 and h10-h11 (Figure 3C) while H3^tail^ segment ^14^KAPRKQLATKAA^25^ makes extensive interactions with h11-h15 (Figure 3B). Density for the segment spanning ^10^STGG^13^ is weak and non-contiguous, consistent with few to no contacts with IMP4. The locations of the N-terminus of H4 and both C-termini of H3 and H4 in the histone-fold domain place these tails far from IMP4, and no density can be observed, suggesting that N-terminal H4^tail^ and the C-terminal tails of H3 and H4 are disordered in the complex and do not contribute to the binding of IMP4 (Figure 3A).

**Figure 3.**
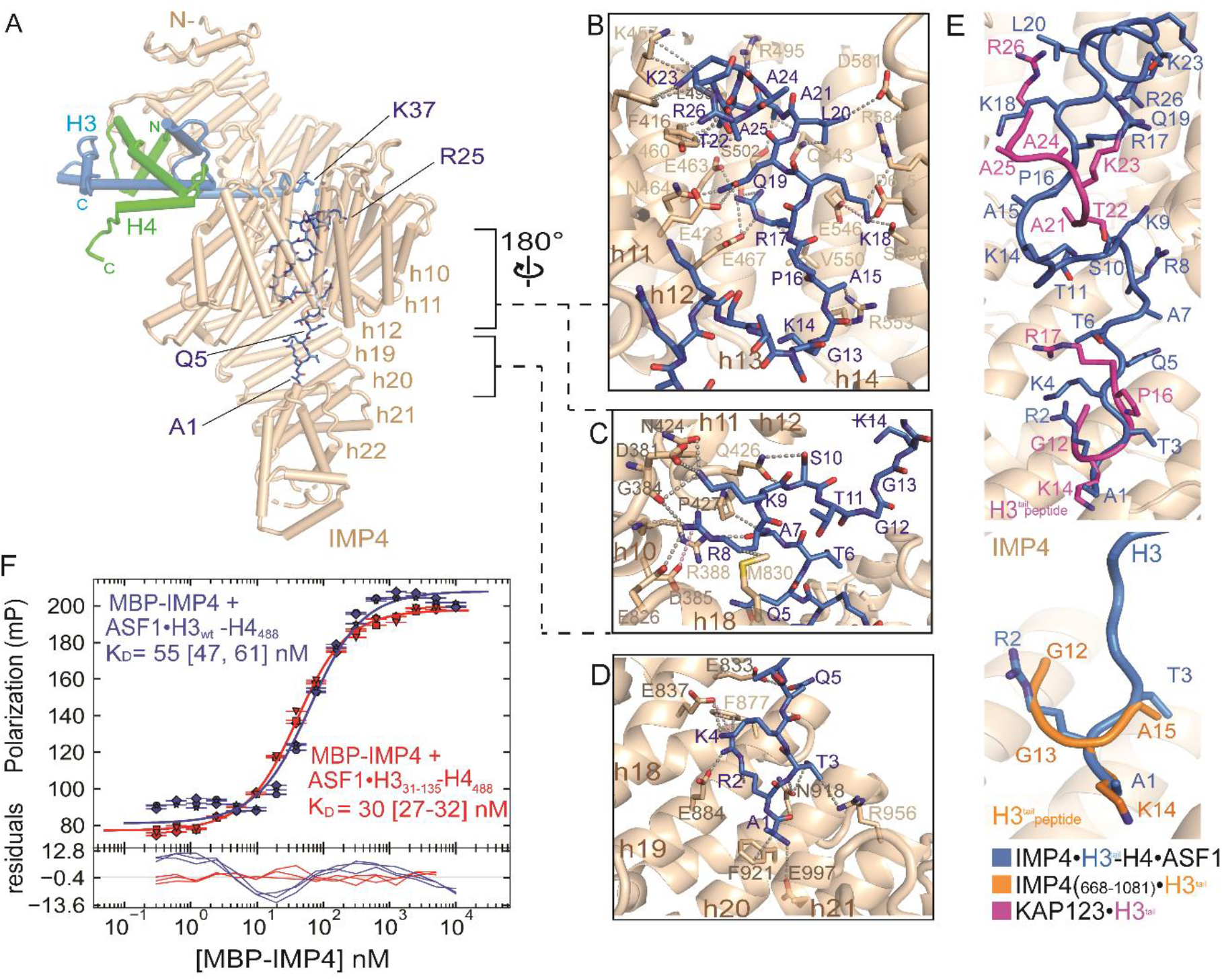
The H3^tail^ binds the C-terminal half of IMP4. A) A view (IMP4 beige, H3 blue, H4 green and ASF1 yellow) showing H3^tail^ residues 1-25 (blue sticks) binding to the C-terminal half of IMP4. B)-D) Details of IMP4-H3^tail^ interactions, which mainly involve side chains (grey dashed lines) and include a few long range electrostatic contacts (pink dashes). B) H3^tail^ residues 14-25 contact residues from HEAT repeats h11-h15 of IMP4; C) H3^tail^ residues 7-10 bind to HEAT repeats h8-h11 and h18; D) H3^tail^ residues 1-5 bind to repeats h18-h22. E) Superposition of H3^tail^ residues of three different structures: IMP4·H3-H4-ASF1 (blue), IMP4_668-1081_·H3_1-18_ (orange; PDBID:5XBK) and KAP123·H3_1-28_ (magenta; PDBID:5VE8). Backbone residues of the H3^tail^ are shown in ribbons representation and the sidechainsare shown in sticks. H3^tail^ IMP4 is beige and in cartoon representation. F) Fluorescence polarization binding assays with MBP-IMP4 and ASF1·H3_WT_-H4_AF488_ (green line) or ASF1·H3_31-135_-H4_AF488_ (blue line). H4 is labeled with the XFD488 fluorophore conjugated to residue 63 which is mutated to cysteine. Fitted binding curves are overlaid onto data points with error bars representing the mean and standard deviation of triplicate titrations. Dissociation constants (*K*_D_s) are reported in the graphs with confidence interval of 68 % in brackets.

Previous biochemical studies had suggested a role for the H3^tail^ in IMP4-binding (Blackwell et al., 2007; Ejlassi et al., 2017; Soniat et al., 2016), but density for the H3^tail^ in our cryo-EM map is not contiguous and weaker than that of the H3-H4 histone-fold domain (Figure S3A-B). We removed the N-terminal tail from H3 (H3Δtail), assembled the wild type (wt) and mutant ASF1·H3-H4 complexes and measured their affinities for IMP4 by fluorescence polarization (Figure 3F and Figure S4). The wt and H3Δtail ASF1·H3-H4 complexes bind IMP4 with similar high affinities (dissociation constants or KDs of 55 nM and 30 nM, respectively), suggesting minor energetic contributions of the H3^tail^ to formation of the IMP4·H3-H4·ASF1 complex.

The H3^tail^ in our structure binds somewhat differently to IMP4 than in previous crystal structures of the H3^tail^ bound to an IMP4 fragment and to the homologous KAP123, respectively. In the IMP4_668-1081_·H3^tail^ structure (5XBK), only four H3^tail^ residues (^12^GGKA^15^) of the 28-residue tail peptide were modeled into a site occupied by H3 residues ^1^ART^3^ in our structure, but in a different orientation (bottom panel, Figure 3E) (An et al., 2017; Soniat and Chook, 2016; Yoon et al., 2018). A slightly longer ^12^GGKAPR^17^ segment was modeled in same site of the KAP123 ·H3^tail^ structure (5VE8) along with another short H3^tail^ segment ^22^SKAAR^27^ binding to KAP123 h11-h13 (An et al., 2017; Soniat and Chook, 2016; Yoon et al., 2018), close to the binding site for H3^tail 14^KAPR^17^ in our structure (top panel, Figure 3E). However, a previous report of the extreme N-terminus of the H3^tail^ (residues K4 and K9) crosslinked with the C-terminal region of IMP4 (Yoon et al., 2018) better supports our model of the IMP4-bound H3^tail^. Our IMP4·H3-H4·ASF1 cryo-EM structure is also different from the integrative model that was built based on crosslinking mass spectrometry and negative-stained EM data (Yoon et al., 2018). IMP4 in the integrative model (coordinates not deposited) seems to adopt a more open conformation and binds mostly to the H3^tail^. Their H3-H4 histone-fold domain·ASF1 was placed close to the C-terminal half of IMP4, in contrast to our ASF1-bound H3-H4 globular domain, which is clamped in the N-terminal half of IMP4.

ASF1 binds the globular domain of H3-H4 but makes no direct contact with IMP4. The histone-chaperone shields the site for H3-H4 tetramerization and likely also stabilizes the otherwise unstable globular fold of the H3-H4 dimer, allowing it to bind IMP4 (Banks and Gloss, 2003; English et al., 2006). IMP4 further shields the H2A-H2B and DNA sites of H3-H4 found in the nucleosome (Figure 4A). Together, IMP4 and ASF1 act in concert as histone-chaperones to shield much of the nucleosome interaction sites of H3-H4. This is yet another example of importins acting as histone-chaperones while transporting these very abundant highly charged histones that are very prone to aggregation; Importin-9 is an effective chaperone of its cargo H2A-H2B and the Importin-7/Importin-β heterodimer effectively shields the linker histone H1 (Ivic et al., 2019; Padavannil et al., 2019).

**Figure 4.**
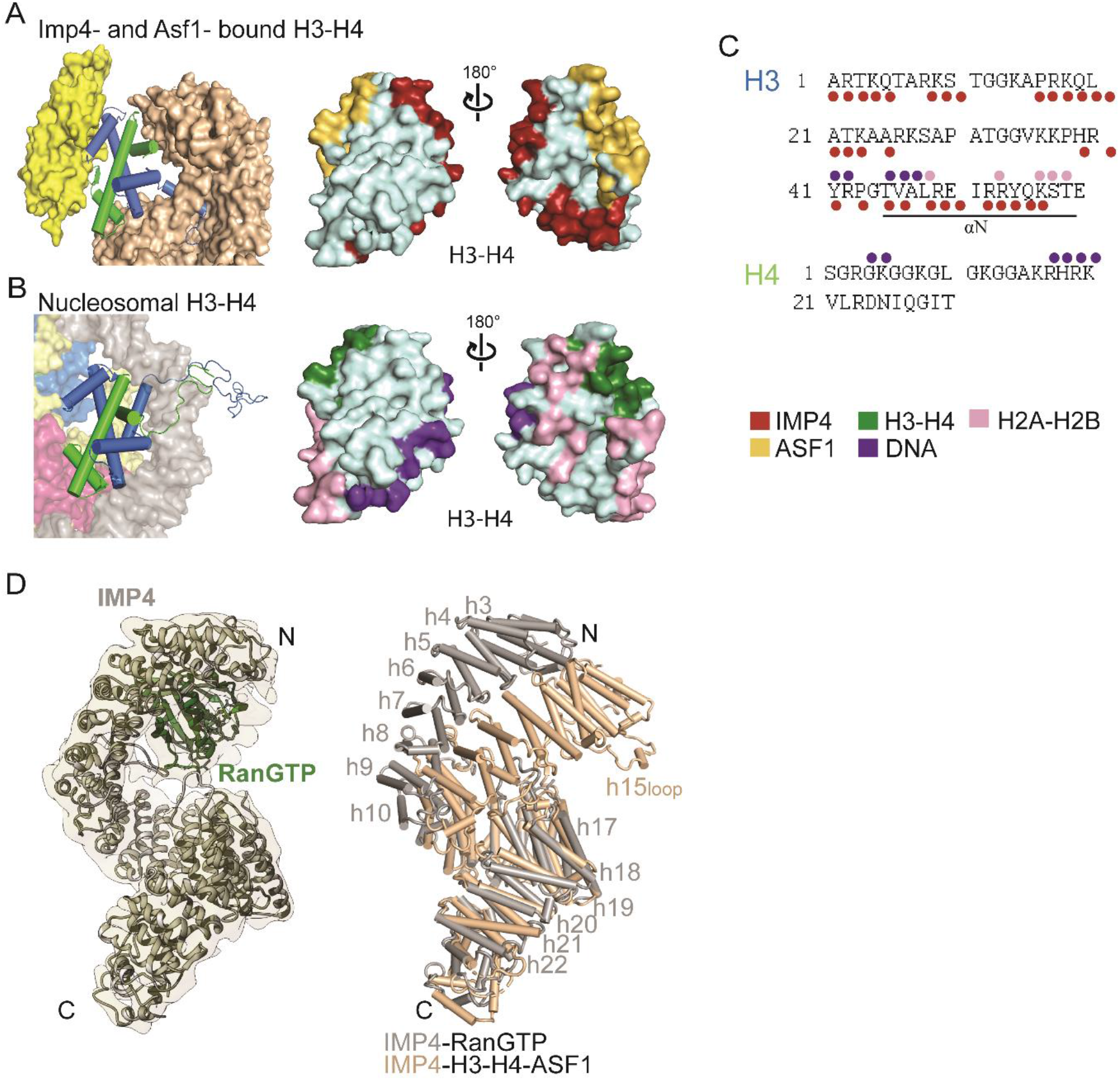
IMP4 and ASF1 act together to shield H3-H4 surfaces and large IMP4 conformational changes occur upon RanGTP-binding. A) Left: H3-H4 dimer (blue-green cartoon) bound to IMP4 (beige surface) and ASF1 (yellow surface). Right: H3-H4 surface with IMP4 interface (red) and ASF1 interface (yellow). B) Left: one H3-H4 dimer (cartoon) in the nucleosome, bound to DNA (grey surface), H2A-H2B (pink-yellow surfaces) and another H3-H4 dimer (blue-green surfaces). Right: H3-H4 surface with the nucleosomal DNA interface (purple) and H2A-H2B interface (pink) and the H3-H4 interface (green). C) H3^tail^ and H4^tail^ sequences: residues that bind IMP4 in the IMP4·H3-H4·ASF1 structure are marked with red circles, residues that contact DNA and H2A-H2B in the nucleosome are marked with purple and pink circles, respectively. D) Left, the 7.1 Å Cryo-EM map of the IMP4·RanGTP complex, with IMP4 (gray cartoon) and RanGTP (dark green cartoon). Right, superposition of repeats h9-h18 of IMP4·H3-H4-ASF1 (IMP4 is beige) and Imp4·RanGTP (IMP4 is grey) showing large conformational change of the N-terminal half of IMP4 relative to its C-terminal half to accommodate RanGTP.

In the nucleus, RanGTP binds Importins with high affinity and dissociates H3-H4·ASF1 from IMP4, ending the nuclear import process (Soniat et al., 2016; Wing et al., 2022). We obtained a 7.1 Å resolution cryo-EM map of the IMP4·RanGTP complex; the HEAT repeat helices of IMP4 can be easily modeled and RanGTP could be reliably docked into the map by aligning the N-terminal HEAT repeats of IMP4 with the Kap121·RanGTP structure (PDBID 3W3Z) (Figure 4B). Alignment of H3-H4·ASF1- and RanGTP-bound IMP4 proteins (Figure 4B) and a Pymol morph (Figure S6) suggest large conformational differences in the IMP4 superhelix. HEAT repeats h2-h8 (rmsds 1.7 Å) and h10-h18 (rmsd 1.4 Å) act as two rigid bodies that move relative to each other about a hinge at h8-h10.

The h15^loop^ also undergoes a large conformational change. In the IMP4·H3-H4·ASF1 structure, h15^loop^ is ordered and bound to both the N-terminal HEAT repeats and the H3-H4 globular domain. However, in IMP4·RanGTP structure, Ran binds to the same IMP4 N-terminal HEAT repeats and displaces the h15^loop^; no density is observed for the loop. The different arrangements of the superhelix and the h15^loop^ result in 1) a smaller N-terminal IMP4 arch that clamps the smaller H3-H4 domain along with a closed and tight central ring that the H3^tail^ threads through in the histone-bound IMP4, *versus* 2) a larger N-terminal IMP4 arch that binds RanGTP in the GTPase-bound IMP4. Rearrangement of the Ran-bound superhelix also separates the central HEAT repeats and opens the central ring. Binding sites for H3-H4 are lost in this major rearrangement of the RanGTP-bound IMP4 superhelix.

In conclusion, the cryo-EM structure of IMP4·H3-H4·ASF1 shows the entire length of the IMP4 making extensive interactions with the H3-H4 dimer, which also binds and is stabilized by ASF1. The N-terminal half of IMP4 binds the H3-H4 histone-fold domain and the αN^H3^ helix, while the C-terminal half of IMP4 binds the H3^tail^. We showed that despite interactions with IMP4, the H3^tail^ is dispensable for formation of the IMP4·H3-H4·ASF1 complex. Finally, large conformational changes of the IMP4 superhelix and the long h15^loop^ allows RanGTP to bind to IMP4 and release H3-H4·ASF1 into the nucleus.

## Resource Availability

### Lead contact and Material Availability

Further information and requests for resources and reagents should be directed to and will be fulfilled by the Lead Contact, Yuh Min Chook.

### Data and code availability

Will provide upon submission to the PDB.

## Experimental model and subject details

### Protein constructs

The gene coding for the full-length human IMP4 was cloned into modified pGEX4t3 and pMALHis6 vectors that contain TEV protease cleavage sites (Chook and Blobel, 1999; Chook et al., 2002). The cDNA fragment corresponding to yeast ASF1(1-160) was cloned into a pMALHis6 vector. Mutant and wt *X. laevis* H3 and H4 were cloned into pET3a and the H4_E63C_ construct was obtained from The Histone Source (Colorado State University). Truncated yeast Ran (residues 1-179) with the Q71L mutation to prevent GTP hydrolysis was cloned in pET21d (Fung et al., 2015).

### Method details

#### Protein expression and purification

##### IMP4

was expressed as MBP-IMP4 or GST-IMP4 in *E. coli* BL21 cells, grown in LB media and induced by adding 0.5 mM of IPTG for 12h at 25°C. Cells were harvested, resuspended in buffer containing 50 mM Tris HCl (pH 7.5), 500 mM NaCl, 2 mM EDTA, 20 % glycerol and protease inhibitors, and lyzed by high pressure cell disruption. After centrifugation of the lysate, the supernatant was collected for purification. The supernatant was incubated with amylose resins (New England BioLabs) or GSH resins (Cytiva), which were later washed several times with buffer containing 50 mM Tris HCl (pH 7.5), 150 mM NaCl and 20 % glycerol. GST-IMP4 was then cleaved with TEV protease (4°C for 12h) to remove the GST-tag, and IMP4 was eluted in the same buffer. MBP-IMP4 was eluted from the amylose resin by adding 30 mM of D-maltose (Sigma) to the elution buffer and the eluted sample was further purified by ion exchange chromatography using HP-Q column followed by size exclusion chromatography using a Superdex 200 Increase 10/300 GL column (Cytiva) that was pre-equilibrated with buffer containing 50 mM Tris HCl (pH 7.5), 500 mM NaCl and 20 % glycerol. Peaks corresponding to the IMP4 or MBP-IMP4 were collected and concentrated for the experiments.

##### ASF1

Supernatant containing ASF1(1-160) was incubated with amylose resin (New England BioLabs), washed with buffer containing 50 mM Tris HCl (pH 7.5), 150 mM NaCl and 20 % glycerol, and then eluted by cleavage with TEV protease. The eluted ASF1 sample was further purified by ion exchange using a HP-Q column and then concentrated for injection onto a size-exclusion column that was pre-equilibrated with buffer containing 30 mM Tris (pH 8.0), 150 Mm NaCl, 10% glycerol.

##### H3-H4

Wt and mutant *X. laevis* H3 and H4 were expressed in *E. coli* BL21 (DE3) pLysS cells by inducing expression with the addition of 0.5 mM of IPTG to the 2xYT media for 4h at 25°C. After lysis by sonication in a buffer containing 50 mM Tris HCl (pH 7.5), 150 mM NaCl, 2 mM EDTA and protease inhibitors, followed by centrifugation, the pellet was washed several times, then resuspended with denaturing buffer (7 M guanidine HCl, 50 mM Tris HCl (pH 7.5) and 10 mM DTT) and dialyzed overnight in SAU-200 buffer (7 M urea, 20 mM sodium acetate (pH 5.2), 200 mM NaCl, 1 mM EDTA, 5 mM □-mercaptoethanol). Unfolded H3 and H4 proteins were purified separately by cation exchange chromatography using a SP column, in SAU buffer with a gradient of 200 to 600 mM NaCl. (H3-H4)2 tetramers were formed by mixing H3 and H4 in an equimolar ratio, followed by dialysis overnight in a buffer without guanidine HCl (50 mM Tris HCl (pH 7.5), 2 M NaCl, 5 mM □-mercaptoethanol). H3-H4 tetramers were purified by size-exclusion chromatography in a Superdex 200 column. The ASF1·H3-H4 complex was formed by mixing 1:1 molar ratio of Asf1(1-160) and (H3-H4)2 tetramers. The ASF1·H3-H4 complex was purified by size-exclusion chromatography, which removed excess Asf1. The complex IMP4·H3-H4·ASF1 was formed by incubating IMP4 and ASF1·H3-H4 in 1:1 molar ratio for 30 min and then purified by size-exclusion chromatography. Fractions were concentrated to 5-10 mg/ml for cryo-EM sample grid preparation.

##### GSP1

Sc Ran (1-179, Q71L) was expressed in *E. coli* BL21 (DE3) by induction with 0.5 mM IPTG for 12h at 20 °C. Cell were harvested and lysed by high pressure cell disruption and the supernatant was incubated with Ni-NTA Agarose resins (QIAGEN) and eluted in buffer containing 50 mM HEPES pH 7.4, 100 mM NaCl, 10 % Glycerol, 2 mM MgOAc_2_, 2 mM DTT, 500 mM Imidazole. To load the GTPase with GTP, RAN was first incubated with 7 mM EDTA for 30 minutes followed by 30 minutes incubation with 20 mM MgOAc_2_ and 7 mM GTP. RANGTP was further purified by cation exchange chromatography using a SP column equilibrated with 20 mM HEPES pH 7.4, 4 mM MgOAc, 1 mM DTT, 10 % Glycerol and eluted with a gradient of 0-1 M NaCl. To form the IMP4-RANGTP complex, proteins were incubated in a 1:1 molar ratio for 30 min and then purified by size-exclusion chromatography.

#### Cryo-EM grid preparation, data collection and data processing

Samples of IMP4·H3-H4·ASF1 and IMP4·RANGTP were buffer exchanged into a final buffer containing 50 mM Tris HCl (pH 7.5), 150 mM NaCl and 0.1 % NP-40 at a final protein concentration of 3 mg/ml. 3.5 μl of the IMP4·H3-H4·ASF1 or IMP4·RANGTP samples were applied on holey carbon grids (Quantifoil R1.2/1.3, 300 mesh copper) that were previously glow-discharged using a PELCO easiGlow unit (Ted Pella) for 60 s at 30 mA, and then frozen using the Vitrobot Mark IV System (Thermo Fisher). The grids with the best particle distribution were submitted to 24h data collection. Datasets of both of IMP4·H3-H4·ASF1 and IMP4·RANGTP complexes used for final data processing were collected at the Cryo Electron Microscopy Facility at UT Southwestern on a Titan Krios microscope (Thermo Fisher) operating at 300 kV, with the post-column energy filter (Gatan) and a K3 direct detection camera (Gatan), using SerialEM (ref. Mastronarde, 2005). For the IMP4·H3-H4·ASF1 complex, 4,126 movies were acquired at a pixel size of 0.415 Å in super-resolution counting mode, with an accumulated total dose of 52 e-/Å2 over 40 frames. The defocus range of the images was set to be −1.0 to −2.5 μm. For the IMP4·RANGTP complex, 2,933 movies were acquired at a pixel size of 0.545 Å in super-resolution counting mode, with an accumulated total dose of 50 e-/Å2 over 50 frames. The defocus range of the images was set to be −1.2 to −2.7 μm.

All data processing was performed using the software cryoSPARC v3.3.1 (Punjani et al., 2017). A ½ F-crop factor was applied during motion correction, followed by patch CTF estimation. A small subset of ~20 frames was used to generate the initial template for particle picking. For the IMP4·H3-H4·ASF1 complex, initially a total of 2,636,349 particles extracted from all the micrographs. 217,815 particles were selected after the initial round of 2D classification. 146,050 particles were included after additional four rounds of 2D classification for *ab initio* modeling followed by heterogeneous refinement. Non-uniform refinement was carried out to generate the final 3.45 Å resolution map. For the IMP4·RANGTP complex, 1,226,438 particles were extracted from all the micrographs. 16,089 particles were selected after 9 rounds of 2D classification and used to generate *ab initio*_models followed by heterogeneous refinement. Homogeneous refinement led to the final 7.1 Å resolution map.

#### Cryo-EM model building, refinement and analysis

##### IMP4·H3-H4·ASF1

The ASF1·H3-H4 (PDBID: 2HUE) and KAP121 (PDBID: 3W3T) structures were used to build the model of ASF1·H3-H4·IMP4. SWISS-MODEL was used to generate a 3D-model of IMP4 using KAP121 as template. First, the models were docked into the cryo-EM map using UCSF Chimera (Pettersen et al., 2004), followed by multiples cycles of manual model building and refinement using Coot (Emsley, 2018). The final model was subjected to real-space refinement in Phenix (Adams et al., 2010). The binding interfaces were then analyzed using CONTACT/ACT (Winn et al., 2011) with a contacts cut-off of 4.0 Å. These interactions were curated and analyzed in PyMOL where the final figures were generated (DeLano, 2009).

##### IMP4·RANGTP

IMP4 was modeled into the cryo-EM map using IMP4 from our IMP4·H3-H4·ASF1 structure. RANGTP was docked into the Cryo-EM map, using the PDB coordinated of KAP121-RANGTP as a template, by aligning the HEAT repeats h1-h4 of KAP121 and IMP4. The models were submitted to cycles of model building and refinement using Coot and Phenix (Adams et al., 2010; Emsley, 2018).

#### Fluorescence polarization assays

Dissociation constants (K_D_ values) of IMP4 binding to H3-H4·ASF1 complexes were obtained by fluorescence polarization assays, previously described by (Fung et al., 2015). For these assays, the H4_E63C_ mutant protein (The Histone Source) was labeled with the XFD488 fluorophore (AAT Bioquest) by mixing H4_E63C_ and the dye at 1:4 molar ratio in the histone unfolding buffer (50 mM Tris HCl pH 7.5 and 5M guanidine HCl) without DTT, for 4h at 25 °C, in the dark. Labeling efficiency was further confirmed by intact mass-spectrometry. H3-H4_E63C_ tetramers, ASF1·H3_wt_-H4_E63C_ and ASF1·H3_31-135_-H4_E63C_ were formed as for the wt complexes. MBP-IMP4 and the pre-assembled ASF1·H3-H4 complexes were separately dialyzed in buffer containing 50 mM Hepes pH 7.5, 300 mM NaCl and 2 mM MgCl_2_. MBP-IMP4 was serially diluted from 10 μM to 0.3 nM in the presence of 40 nM of either ASF1 ·H3_wt_-H4_E63C_ or ASF1·H3_31-135_-H4_E63C_ in a 384-well microplate (Corning). Fluorescence signal for each titration was measured using an Data were processed in PALMIST (Scheuermann et al., 2016) using averages of triplicate experiments and the final figures were generated with GUSSI (Brautigam, 2015).. Data were processed in PALMIST (Scheuermann et al., 2016) using averages of triplicate experiments and the final figures were generated with GUSSI (Brautigam, 2015).

## Acknowledgments

We thank the Structural Biology Laboratory and the Cryo-EM Facility at UTSW, which are partially supported by grant RP170644 from the Cancer Prevention & Research Institute of Texas (CPRIT), for cryo-EM studies for their assistance with cryo-EM data collection. We also thank the Erzenberger lab for using their equipment for the plate reader. This work was funded by NIGMS of NIH under Awards R35GM141461 (Y.M.C.), R01GM069909 (Y.M.C.), the Welch Foundation Grants I-1532 (Y.M.C.), support from the Alfred and Mabel Gilman Chair in Molecular Pharmacology, Eugene McDermott Scholar in Biomedical Research (Y.M.C.), Mary Kay International Postdoctoral Fellowship (N.E.B.) and the Gilman Special Opportunities Award (H.Y.J.F.).

## Author contributions

Y.M.C. conceptualized and designed the project. N.E.B. planned and performed the experiments. H.Y.J.F. conceptualized the fluorescence polarization experiments. Z.C. and Y.L. collected the cryo-electron microscopy data and provided advice on data processing. Y.M.C, N.E.B. and H.Y.J.F. analyzed the data. Y.M.C. and N.E.B. wrote the manuscript.

## References

Adams, P.D., Afonine, P.V., Bunkoczi, G., Chen, V.B., Davis, I.W., Echols, N., Headd, J.J., Hung, L.W., Kapral, G.J., Grosse-Kunstleve, R.W., et al. (2010). PHENIX: a comprehensive Python-based system for macromolecular structure solution. Acta Crystallogr D 66, 213–221.

Alvarez, F., Munoz, F., Schilcher, P., Imhof, A., Almouzni, G., and Loyola, A. (2011). Sequential establishment of marks on soluble histones H3 and H4. J Biol Chem 286, 17714–17721.

An, S., Yoon, J., Kim, H., Song, J.J., and Cho, U.S. (2017). Structure-based nuclear import mechanism of histones H3 and H4 mediated by Kap123. Elife 6.

Apta-Smith, M.J., Hernandez-Fernaud, J.R., and Bowman, A.J. (2018). Evidence for the nuclear import of histones H3.1 and H4 as monomers. EMBO J 37.

Baake, M., Doenecke, D., and Albig, W. (2001). Characterisation of nuclear localisation signals of the four human core histones. J Cell Biochem 81, 333–346.

Banks, D.D., and Gloss, L.M. (2003). Equilibrium folding of the core histones: the H3-H4 tetramer is less stable than the H2A-H2B dimer. Biochemistry 42, 6827–6839.

Benson, L.J., Gu, Y., Yakovleva, T., Tong, K., Barrows, C., Strack, C.L., Cook, R.G., Mizzen, C.A., and Annunziato, A.T. (2006). Modifications of H3 and H4 during chromatin replication, nucleosome assembly, and histone exchange. J Biol Chem 281, 9287–9296.

Blackwell, J.S., Jr., Wilkinson, S.T., Mosammaparast, N., and Pemberton, L.F. (2007). Mutational analysis of H3 and H4 N termini reveals distinct roles in nuclear import. J Biol Chem 282, 20142–20150.

Brautigam, C.A. (2015). Calculations and Publication-Quality Illustrations for Analytical Ultracentrifugation Data. Method Enzymol 562, 109–133.

Campos, E.I., Fillingham, J., Li, G., Zheng, H., Voigt, P., Kuo, W.H., Seepany, H., Gao, Z., Day, L.A., Greenblatt, J.F., et al. (2010). The program for processing newly synthesized histones H3.1 and H4. Nat Struct Mol Biol 17, 1343–1351.

Chook, Y.M., and Blobel, G. (1999). Structure of the nuclear transport complex karyopherin-beta2-Ran x GppNHp. Nature 399, 230–237.

Chook, Y.M., Jung, A., Rosen, M.K., and Blobel, G. (2002). Uncoupling Kapbeta2 substrate dissociation and ran binding. Biochemistry 41, 6955–6966.

DeLano, W.L. (2009). PyMOL molecular viewer: Updates and refinements. Abstr Pap Am Chem S 238.

Ejlassi, A., Menil-Philippot, V., Galvani, A., and Thiriet, C. (2017). Histones H3 and H4 require their relevant amino-tails for efficient nuclear import and replication-coupled chromatin assembly in vivo. Sci Rep 7, 3050.

Emsley, P. (2018). New Tools for Ligand Refinement and Validation in Coot and CCP4. Acta Crystallogr A 74, A390–A390.

English, C.M., Adkins, M.W., Carson, J.J., Churchill, M.E., and Tyler, J.K. (2006). Structural basis for the histone chaperone activity of Asf1. Cell 127, 495–508.

Fung, H.Y., Fu, S.C., Brautigam, C.A., and Chook, Y.M. (2015). Structural determinants of nuclear export signal orientation in binding to exportin CRM1. Elife 4.

Ivic, N., Potocnjak, M., Solis-Mezarino, V., Herzog, F., Bilokapic, S., and Halic, M. (2019). Fuzzy Interactions Form and Shape the Histone Transport Complex. Mol Cell 73, 1191–1203 e1196.

Jasencakova, Z., Scharf, A.N., Ask, K., Corpet, A., Imhof, A., Almouzni, G., and Groth, A. (2010). Replication stress interferes with histone recycling and predeposition marking of new histones. Mol Cell 37, 736–743.

Luger, K., Mader, A.W., Richmond, R.K., Sargent, D.F., and Richmond, T.J. (1997). Crystal structure of the nucleosome core particle at 2.8 A resolution. Nature 389, 251–260.

Mosammaparast, N., Guo, Y., Shabanowitz, J., Hunt, D.F., and Pemberton, L.F. (2002). Pathways mediating the nuclear import of histones H3 and H4 in yeast. J Biol Chem 277, 862–868.

Muhlhausser, P., Muller, E.C., Otto, A., and Kutay, U. (2001). Multiple pathways contribute to nuclear import of core histones. EMBO Rep 2, 690–696.

Padavannil, A., Sarkar, P., Kim, S.J., Cagatay, T., Jiou, J., Brautigam, C.A., Tomchick, D.R., Sali, A., D’Arcy, S., and Chook, Y.M. (2019). Importin-9 wraps around the H2A-H2B core to act as nuclear importer and histone chaperone. Elife 8.

Pardal, A.J., Fernandes-Duarte, F., and Bowman, A.J. (2019). The histone chaperoning pathway: from ribosome to nucleosome. Essays Biochem 63, 29–43.

Pettersen, E.F., Goddard, T.D., Huang, C.C., Couch, G.S., Greenblatt, D.M., Meng, E.C., and Ferrin, T.E. (2004). UCSF chimera - A visualization system for exploratory research and analysis. J Comput Chem 25, 1605–1612.

Punjani, A., Rubinstein, J.L., Fleet, D.J., and Brubaker, M.A. (2017). cryoSPARC: algorithms for rapid unsupervised cryo-EM structure determination. Nat Methods 14, 290-+.

Scheuermann, T.H., Padrick, S.B., Gardner, K.H., and Brautigam, C.A. (2016). On the acquisition and analysis of microscale thermophoresis data. Anal Biochem 496, 79–93.

Schwamborn, K., Albig, W., and Doenecke, D. (1998). The histone H1(0) contains multiple sequence elements for nuclear targeting. Exp Cell Res 244, 206–217.

Soniat, M., Cagatay, T., and Chook, Y.M. (2016). Recognition Elements in the Histone H3 and H4 Tails for Seven Different Importins. J Biol Chem 291, 21171–21183.

Soniat, M., and Chook, Y.M. (2016). Karyopherin-beta2 Recognition of a PY-NLS Variant that Lacks the Proline-Tyrosine Motif. Structure 24, 1802–1809.

Tagami, H., Ray-Gallet, D., Almouzni, G., and Nakatani, Y. (2004). Histone H3.1 and H3.3 complexes mediate nucleosome assembly pathways dependent or independent of DNA synthesis. Cell 116, 51–61.

Wing, C.E., Fung, H.Y.J., and Chook, Y.M. (2022). Karyopherin-mediated nucleocytoplasmic transport. Nat Rev Mol Cell Biol.

Winn, M.D., Ballard, C.C., Cowtan, K.D., Dodson, E.J., Emsley, P., Evans, P.R., Keegan, R.M., Krissinel, E.B., Leslie, A.G., McCoy, A., et al. (2011). Overview of the CCP4 suite and current developments. Acta Crystallogr D Biol Crystallogr 67, 235–242.

Yoon, J., Kim, S.J., An, S., Cho, S., Leitner, A., Jung, T., Aebersold, R., Hebert, H., Cho, U.S., and Song, J.J. (2018). Integrative Structural Investigation on the Architecture of Human Importin4_Histone H3/H4_Asf1a Complex and Its Histone H3 Tail Binding. J Mol Biol 430, 822–841.

