## Supplemental Files for "Structure of Importin-4 bound to the H3-H4·ASF1 histone·histone chaperone complex"

**Supplementary Information**

Table S1. Statistics of Cryo-EM data collection, processing and refinement for IMP4·H3-H4·ASF1 and IMP4·RanGTP.

| | IMP4·H3-H4·ASF1 | IMP4·RanGTP |
|---|---|---|
| **Data collection and processing** | | |
| Magnification | 105,000 | 81,000 |
| Voltage (kV) | 300 | 300 |
| Electron exposure ($e^{-}/Å^2$) | 52 | 50 |
| Defocus range (μm) | -2.5 to -1.0 | -1.2 to -2.7 |
| Pixel size (Å) | 0.83 | 1.09 |
| Symmetry imposed | C1 | C1 |
| Initial particle images (no.) | 2,636,349 | 1,226,438 |
| Final particle images (no.) | 146,050 | 16,089 |
| Map resolution (Å) | 3.5 | 7.1 |
| FSC threshold | 0.143 | 0.143 |
| **Refinement** | | |
| Initial model used (PDBID) | 2HUE, 3W3T | 3W3Z |
| Model resolution (Å) | 2.9/3.2/3.6 | 6.4/6.9/8.0 |
| FSC threshold | 0/0.143/0.5 | 0/0.143/0.5 |
| Map sharpening *B* factor ($Å^2$) | -140 | -657 |
| Model composition | | |
| Nonhydrogen atoms | 11,014 | 9,296 |
| Protein residues | 1401 | 1196 |
| R.m.s. deviations | | |
| Bond lengths (Å) | 0.003 | 0.003 |
| Bond angles (°) | 0.644 | 0.872 |
| **Validation** | | |
| MolProbity score | 1.59 | 2.31 |
| Clashscore | 7.59 | 19.52 |
| Poor rotamers (%) | 0.00 | 0.00 |
| Ramachandran plot | | |
| Favored (%) | 97.03 | 90.82 |
| Allowed (%) | 2.97 | 8.75 |
| Disallowed (%) | 0.00 | 0.42 |

*Supplementary 1*

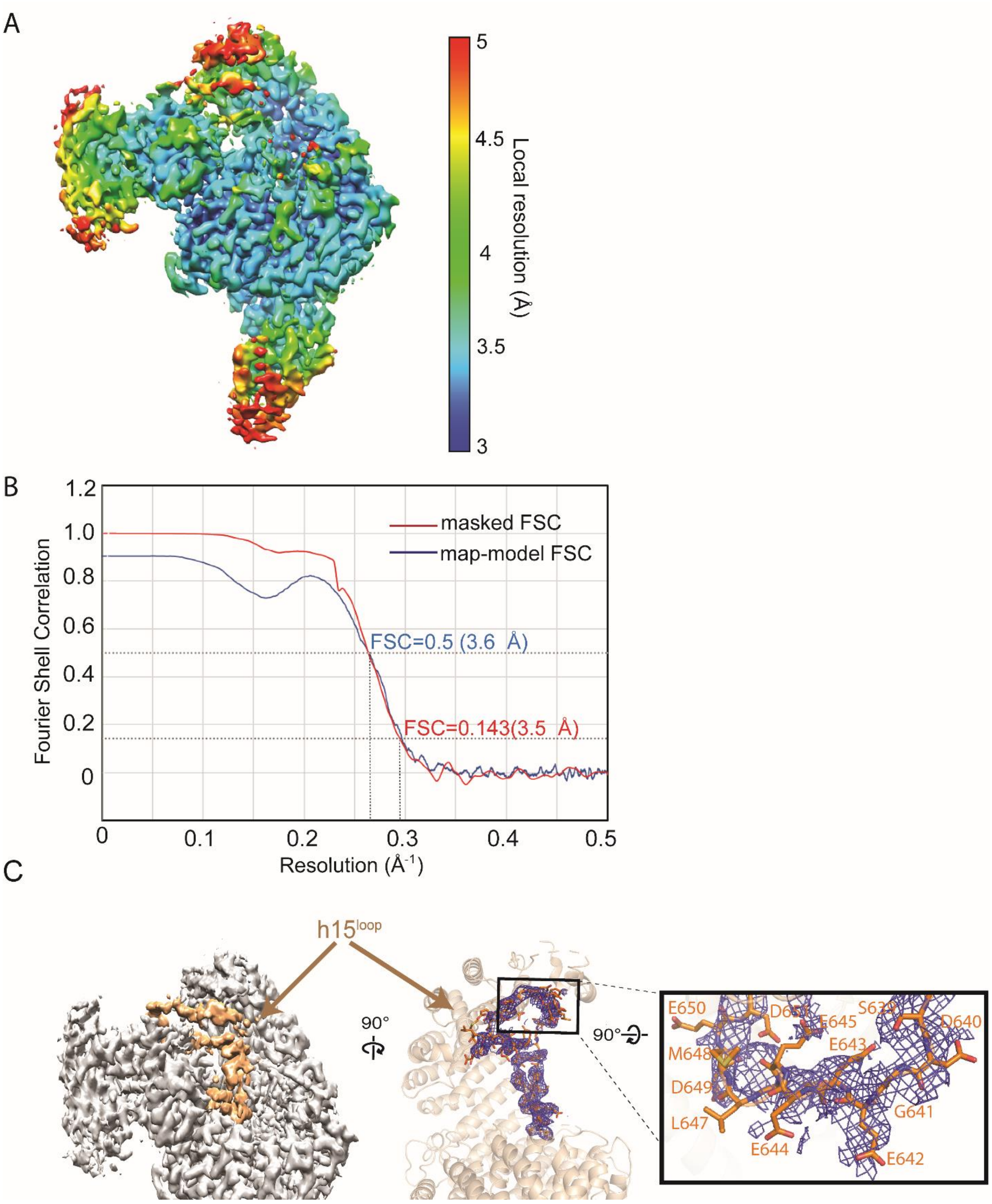


**Figure S1**. **Cryo-EM map of IMP4·H3-H4·ASF1** and density of an IMP4 loop. A) Cryo-EM map colored by local resolution as indicated by the color key. B) The Fourier Shell Correlation (FSC) for the EM density map calculated from cryoSPARC (red line; masked FSC) and phenix.refine (blue line; map-model FSC). C) Left, Cryo-EM map showing the h15$^{loop}$ in orange. Middle, the density of the h15$^{loop}$ shown as blue mesh' contour level = 4 σ. Right, details of the density at h15$^{loop}$ residues 639-651.

*Supplementary* 2

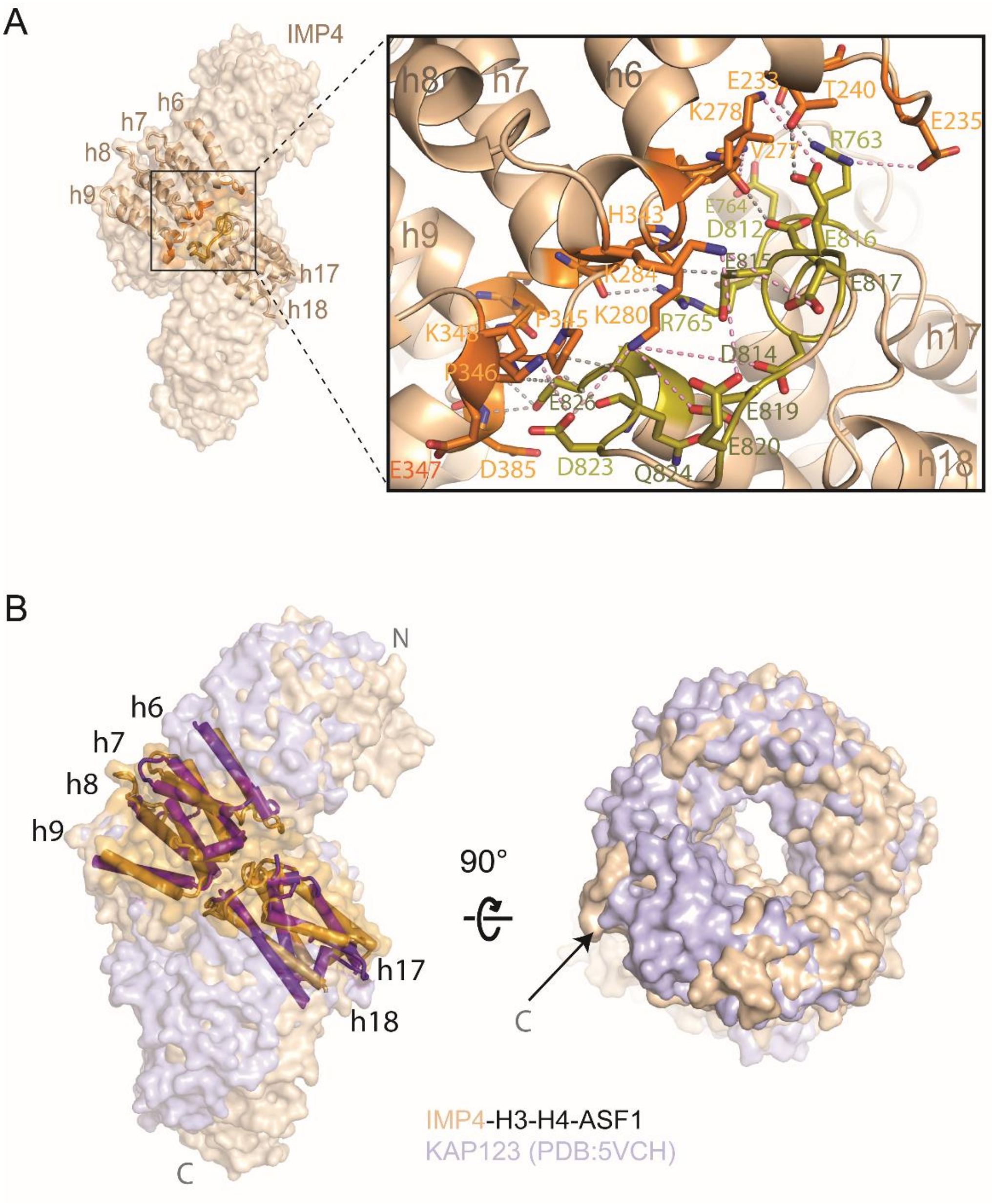


**Figure S2**. **Details of interactions between h6-h9 and h17-h18.** A) Details of the interactions between repeats h6-h9 (orange) and h17-18 (yellow) of IMP4 (beige). Hydrophobic and hydrogen-bonds interactions are shown with gray dashed lines and electrostatic interactions with pink dashes. B) Surface representations of unliganded KAP123 (lilac; PDBID 5VCH) superimposed on the ASF1·H3-H4-bound IMP4 (beige). The helices of repeats h6-h9 and h17-18 are shown as cylinders to highlight the similarities of both importins in this region. The right panel is a 90 ° rotation about the horizontal axis of the left panel to show the central rings of unliganded KAP123 and ASF1·H3-H4-bound IMP4.

*Supplementary 3*

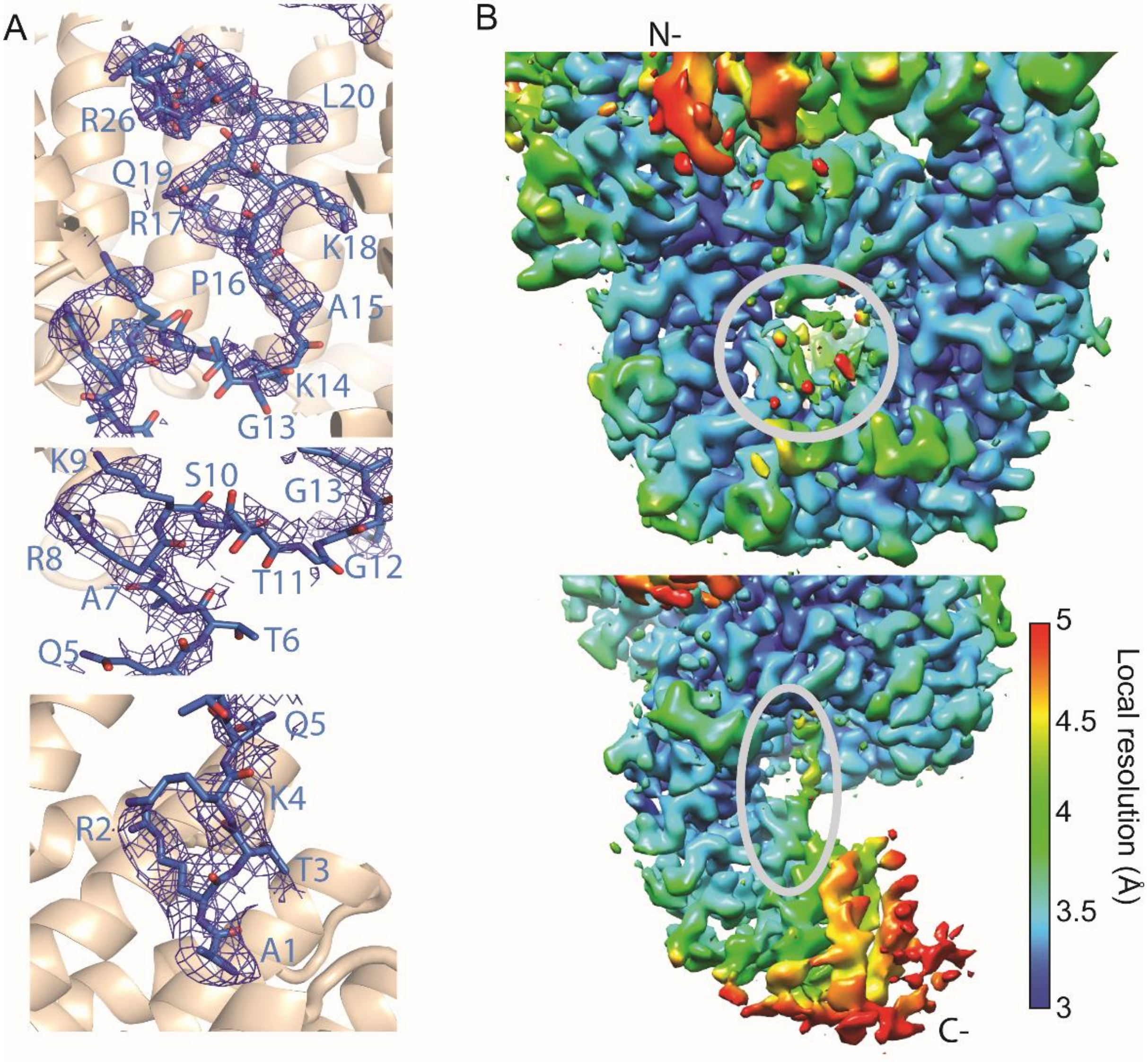


**Figure S3. EM density of the H3$^{tail}$.** A) Cryo-EM density for the H3$^{tail}$ of IMP4·H3-H4·ASF1 (blue mesh; contour level = 4 σ). The H3$^{tail}$ is shown in the same views as in Figure 3B-D. B) EM density of IMP4·H3-H4·ASF1 colored by the local resolution. Top panel, a view of the IMP4 central ring with the H3$^{tail}$ indicated by a light grey circle. Bottom panel, the C-terminal half of IMP4 with the light grey oval highlighting the N-terminal most residues of the H3$^{tail}$.

Supplementary 4

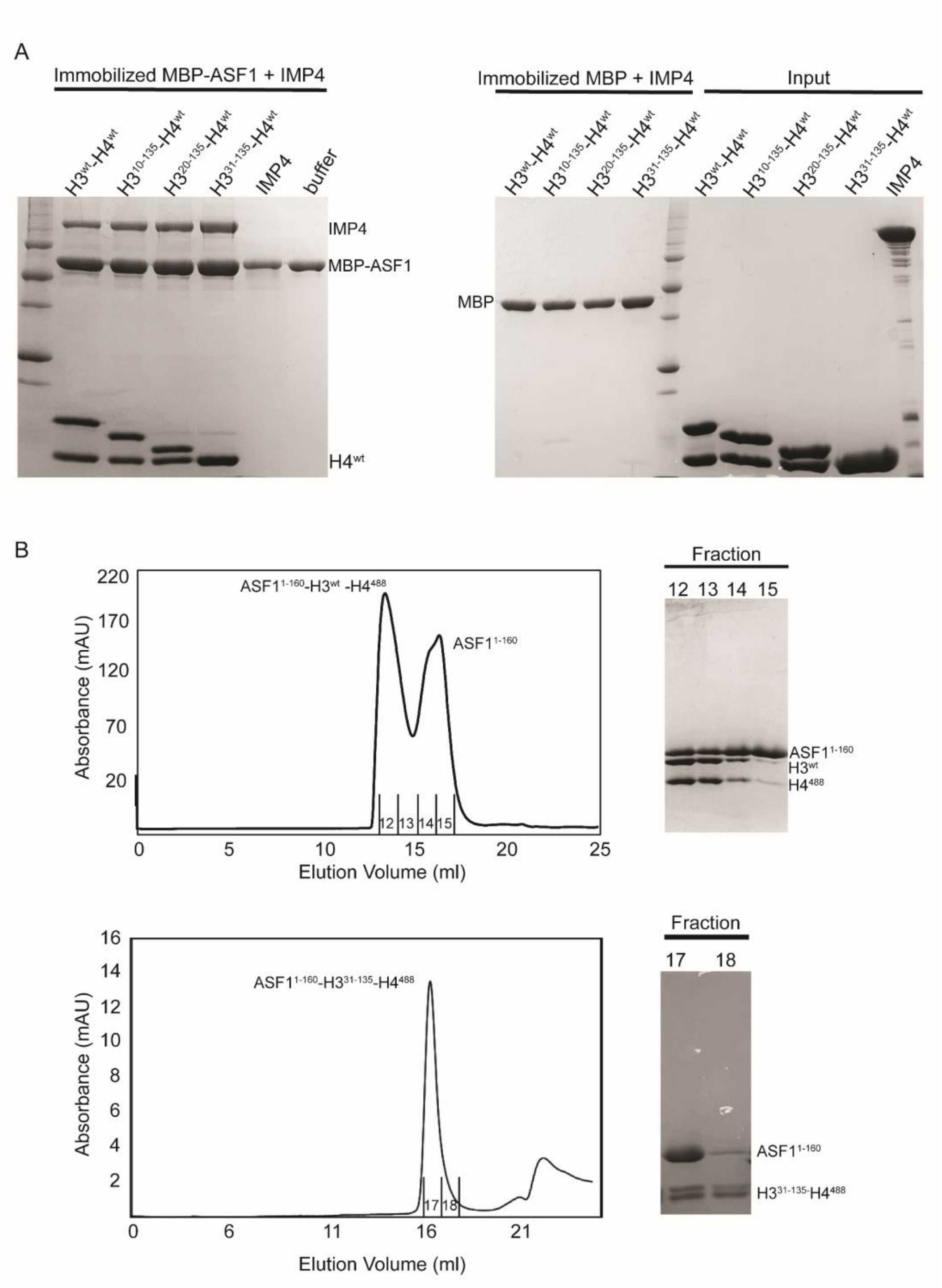


**Figure S4. Binding assays and size-exclusion chromatography of ASF1·H3-H4 with truncated $H3^{tail}$.** A) Pulldown binding assays of immobilized MBP-ASF1 with IMP4 and H3-H4 constructs (SDS-PAGE/Coomassie Blue). B) Size-exclusion chromatography of the ASF1·$H3^{WT}$-$H4^{488}$ and ASF1·$H3^{30\text{-}135}$-$H4^{488}$ complexes used in the fluorescence polarization assays in Figure 3F, and SDS-PAGE of the fractions containing the complexes.

*Supplementary 5*

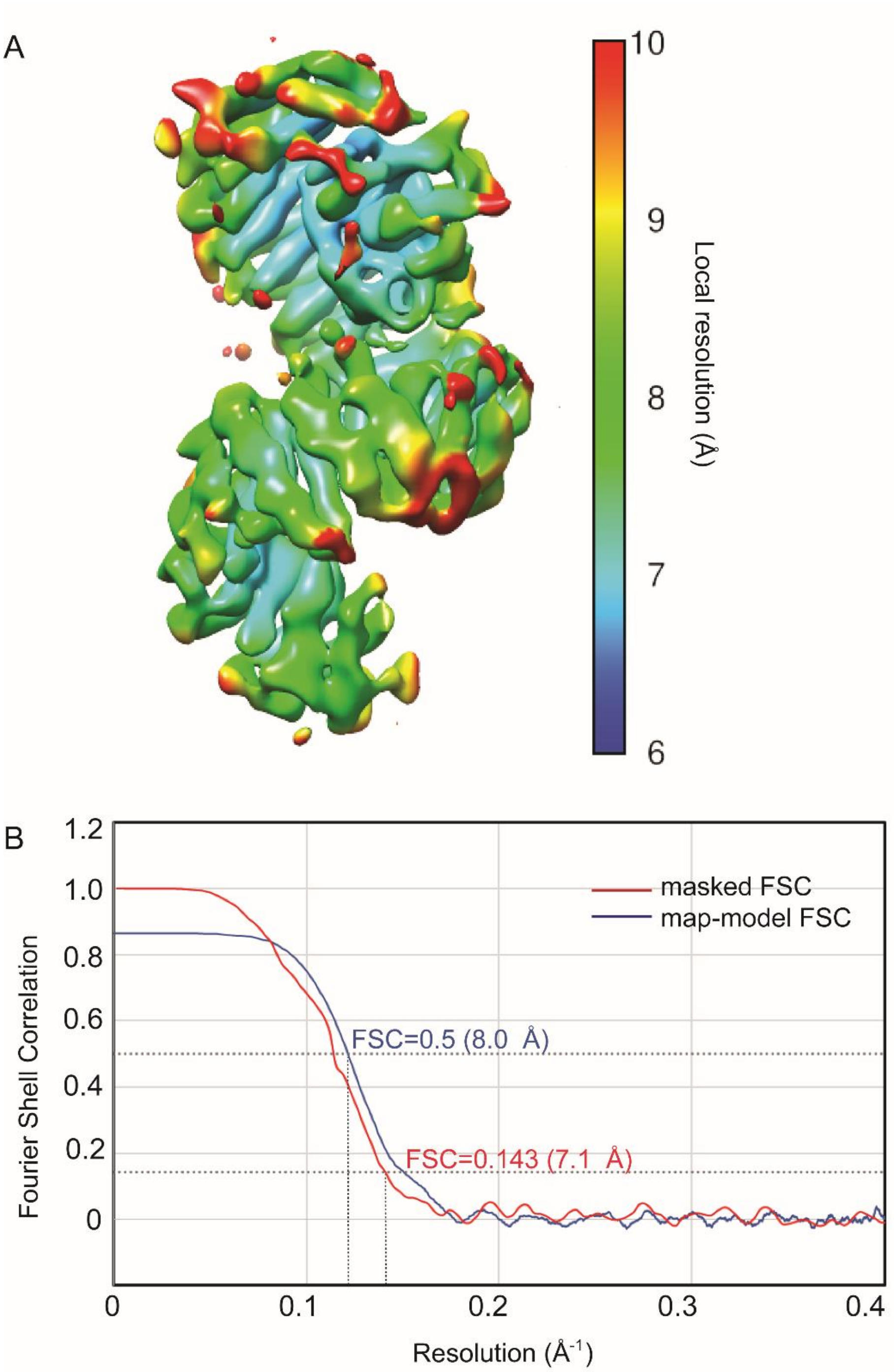


**Figure S5**. **Cryo-EM map of IMP4·RanGTP**. A) Cryo-EM map colored by local resolution as indicated by the color key. B) The Fourier Shell Correlation (FSC) for the EM density map calculated from cryoSPARC (red; masked FSC) and phenix.refine (blue; map-model FSC).
